# Quantifying and understanding reproductive allocation schedules in plants: a lifetime of decisions

**DOI:** 10.1101/008508

**Authors:** Elizabeth Hedi Wenk, Daniel Falster

## Abstract

1. A plant's reproductive allocation (RA) schedule describes the fraction of surplus energy allocated to reproduction as it increases in size. RA schedules distinguish between energy allocated to different tissue types and thus links to a key physiological trade-off in an organism's functioning and life history. While theorists have adopted RA schedules as an elegant connection between life history and energy allocation, little is known about RA schedules in real vegetation.
2. Here we review what is known about RA schedules for woody plants using studies either directly quantifying RA, or which collected data from which the shape of an RA schedule can be inferred.
3. We find that RA schedules vary considerably across species: some species abruptly shift all resources from growth to reproduction (the “big-bang” strategy); most others gradually shift resources into reproduction, but under a variety of graded schedules (“partial bang”, “asymptotic”, “gradual”, and “declining”). Available data suggest the maximum fraction of energy allocated to production ranges from 0.1 to 1 and that shorter-lived species have higher initial RA and increase their RA more quickly than do longer-lived species.
4. ‘*Synthesis*’ Available data suggests a wide variety of RA schedules exist among plant species. The collection of more data on RA, and not only on easy-to-measure proxies such as maximum height, would enable a tighter integration between theory and observation in plant ecology.

## Introduction

A primary goal of plant ecophysiological theory is to break down plant function into a common set of processes that identify strategic differences among individuals and species. By documenting links between individual tissues or allocation decisions on carbon uptake, growth and mortality, plant ecology has moved decidedly towards a trait-centric understanding of vegetation over the last twenty years (Reich, Walters & Ellsworth 1992; Westoby *et al.* 2002; Cornelissen *et al.* 2003; McGill *et al.* 2006; Chave *et al.* 2009; Wright *et al.* 2010). Given a common set of physiological rules describing plant construction and function, differences in growth strategy among species can increasingly be captured via a select number of functional traits (e.g. Falster *et al.* 2011). There is strong evidence for trade-offs associated with leaf functioning, stem construction, plant hydraulics, and the division of reproductive effort into few large or many small seeds (Henery & Westoby 2001; Wright *et al.* 2004; Chave *et al.* 2009; Poorter *et al.* 2010). There also exists substantial and well-documented variation among species in each of these traits (Westoby *et al.* 2002). However, we have a limited understanding of how species differ from one another in the amount of energy they allocate to reproduction.

The partitioning of energy between reproduction and other activities throughout a plant's lifetime – such as growth, storage and defence – is arguably the most fundamental component of its life-history (Harper & Ogden 1970; Bazzaz, Ackerly & Reekie 2000). Here we refer to the fraction of surplus photosynthetic energy that is allocated to reproduction in a given period as reproductive allocation (RA), where surplus energy is that which remains after the costs of respiration and tissue turnover have been paid. The change in RA with respect to size or age will be termed an RA schedule. Figure 1 shows several possible RA schedules. The two most extreme RA schedules include a slow increase in RA across a plant's lifetime (a graded RA schedule) and an RA schedule where maximum RA is reached and vegetative growth ceases as soon as reproduction commences (a big bang schedule, indicating an instantaneous switch from RA=0 to RA=1) (Figure 1). Big-bang reproducers are also termed semelparous or monocarpic, a group that includes some annuals, several succulent shrubs and even a few large trees (Young 2010; Thomas 2011) (Figure 1, panel b). A graded RA schedule, also termed iteroparous or polycarpic, can be further divided into RA schedules we term partial bang, asymptotic, gradual, and declining depending on how RA shifts as the plants age (Figure 1, panels c-g). A plant's RA schedule directly influences its life history via age at first reproduction, fecundity over time, and growth rates. At the simplest level, the choice of allocating energy to growth versus reproduction is between early reproductive onset and high reproductive investment, resulting in the rapid production of abundant seeds at the expense of growth and survival, versus a long period devoted entirely to growth followed by more modest reproductive output, as happens with most large trees. From the perspective of other organisms, the RA schedule determines how gross primary productivity is allocated among fundamentally different tissue types, i.e. leaves, woody tissues, flowers, fruits and seeds, the eventual food stuffs at the base of terrestrial food webs. Figure 2 highlights, using a simple plant growth model from Falster et al 2011, how differences in RA schedule alone can drive differences in growth, seed production and biomass allocation. For this reason, theorists have adopted RA schedules as an elegant connection between life history and energy allocation. Optimal energy models illustrate how RA schedules should respond to a variety of biotic and abiotic factors including tissue turnover, seed set, age-specific mortality, and environmental stochasticity, suggesting substantial variation in RA schedules is to be expected across species with different life histories (Myers & Doyle 1983; Reekie & Avila-Sakar 2005; Miller, Tenhumberg & Louda 2008; Miller *et al.* 2012).

**Figure 1.**
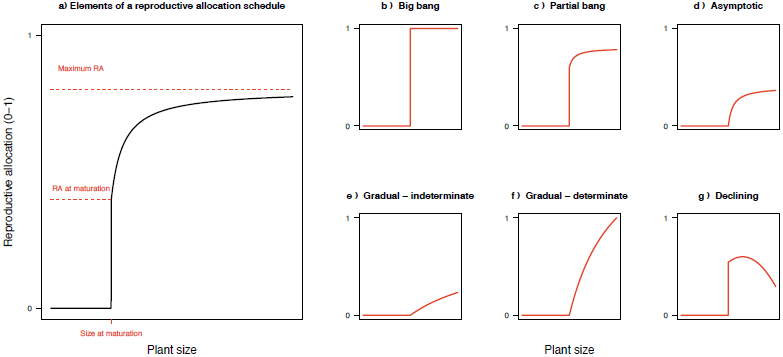
Classifying reproductive allocation schedules. Panel **a** highlights elements of a schedule that can be quantified in their own right, while panels **b-g** illustrate alternative schedules.

**Figure 2.**
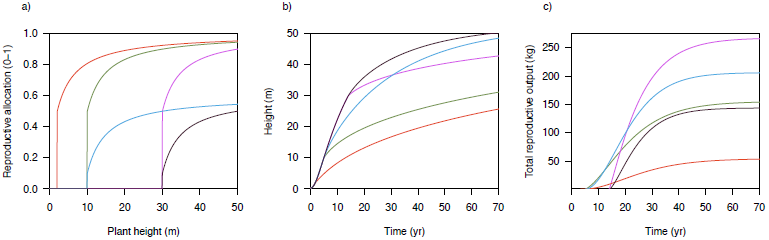
Reproductive allocation schedules influence growth rate, size and seed output. Using a generic model of plant growth (Falster et al 2011), we simulated growth of five individual plants with different RA schedules. Panels b-c show how differences in height and lifetime reproductive output accumulate over time. Full details on model given in Supplementary material.

While RA schedules provide an elegant lifelong summary of the outcome of a plant’s strategy for reproduction, widespread collection of empirical data has been limited due to the effort required to accurately determine the many sinks for surplus energy, including growth, storage, defence and reproduction. In particular, very few data on lifetime reproductive allocation exist for long-lived species, due to the impracticalities of assessing reproductive output across an individual tree’s lifetime. Not surprisingly, research has focussed on other features of reproductive function that are easier to quantify, including instantaneous measures of reproductive output (RO; Henery & Westoby 2001; Weiner *et al.* 2009), plots of reproductive output versus vegetative mass (RV curves; Weiner *et al.* 2009), a species’ maximum height (Wright *et al.* 2010; Cornwell *et al.* 2014), and relative size at onset of maturity (RSOM; Thomas 2011). Below we identify the shortcomings of these metrics in quantifying reproductive patterns and state the case as to why explicitly quantifying RA schedules is needed.

Reproductive output is the instantaneous measure of seed production per unit time (either in numbers or units mass per time). RO is occasionally regarded as a species ‘trait’, yet unlike RA it is not the direct outcome of a trade-off decision. The RO of a plant is the result of the combined effects of its instantaneous RA and its productive capacity, such that two plants of a given size could have identical RO, but one would have higher productive capacity and a lower RA and a second plant could have the reverse. It is well known that RO is also influenced by both a variety of stem and leaf traits and abiotic factors such as resource availability. Moreover, both RA and productive capacity can independently evolve as a plant ages. For instance, RO increases with plant size, for the productive capacity of a plant increases along with its total leaf area (Niklas & Enquist 2003; Falster, Moles, & Westoby 2008; Weiner *et al.* 2009; Figure 3). Meanwhile, a species’ photosynthetic capacity may decrease with age, as has been shown, decreasing its pool of surplus energy (Niinemets 2002; Thomas 2010). The balance of these independence processes means RA itself may either increase or decrease as a plant ages.

**Figure 3.**
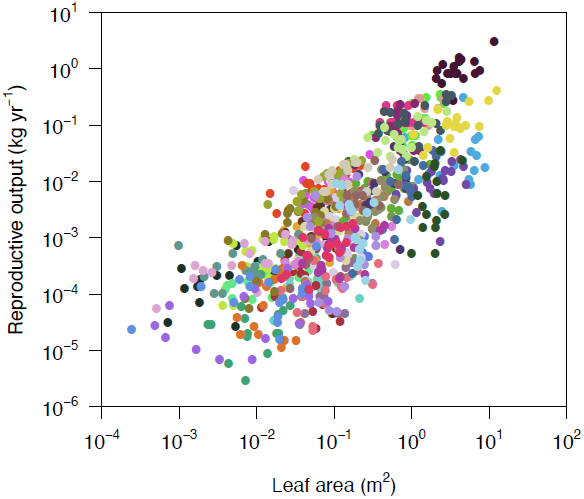
Variation in reproductive output with size within populations for 47 co-occurring species. Data are from (Henery & Westoby 2001). Fruiting and seed production data were collected for 47 woody perennial species over a period of one year in Ku-ring-gai Chase National Park, Australia. In each species, annual fruit production data for six randomly selected reproductively mature individuals per species at each site were collected over a period of 12 months as the fruit matured. Each dot represents an individual; species are distinguished by colours.

Despite these drawbacks, the relationship between plant size and RO is often examined by constructing a log-log regression of cumulative lifetime RO against vegetative size - a so called ‘RV curve’ (Weiner *et al.* 2009). This approach has been used extensively for annuals and herbs, but is difficult to apply to large, long-lived species where lifetime RO is unknown (Weiner *et al.* 2009). Instead of measuring lifetime RO for long-lived organisms, one might instead quantify annual RO, as in Figure 3. Figure 3 presents data on annual RO in relation to size for 47 coexisting plant species. It shows that for most species, RO increases with size but that species differ by at least two orders of magnitude in the amount of production at any given size. Do such differences reflect different levels of photosynthetic productivity? Or do they indicate different levels of allocation to seed production? If one knew the plant’s RA schedule, one could separate the effects of instantaneous RA and productive capacity on RO. An additional drawback of RV curves, is that current vegetative size is not necessarily well correlated with current surplus energy. Vegetative size is indicative of both past and current growth, as well as tissue turnover rates, while its current RO is a function of its instantaneous (or the current seasons) surplus energy. As plants age their surplus energy pool may begin to plateau or even decrease, both through declining photosynthetic capacity and increasing tissue replacement costs. An RV curve cannot distinguish between lower RO due to lower RA or due to decreasing energy budgets. Overall, neither RV curves nor instantaneous RO can be incorporated into process-based vegetation models, since neither incorporates traits that are underpinned by a single trade-off. They therefore have limited utility for understanding how energy is partitioned between growth and reproduction across the lifetime of long-lived organisms.

Maximum height and RSOM are two other metrics used to assess the trade-off between growth and reproduction. The premise for using maximum height is that a species with a greater maximum height has delayed diverting energy to reproduction for longer and hence maintained a greater growth rate for longer during development (Turner 2001; Westoby *et al.* 2002). Supporting this hypothesis, greater maximum height was correlated with higher potential growth rate in tropical forests (Wright *et al.* 2010). The advantage of using maximum height as a proxy for reproductive allocation is that it is easy to measure: data now exist for over 20000 species (Cornwell *et al.* 2014). The main problem with maximum height is that it quantifies the outcome of both demographic luck and a whole host of individual trade-offs, not just the RA trade-off. Moreover, the nature of all these trade-offs may shift with age and / or across its geographic range. As is shown in Figure 2, different RA schedules can yield the same final maximum height, but with different growth rates along the way, leading to different competitive interactions. RSOM, the ratio of threshold size (size at reproductive onset) to maximum size, is likewise used to summarize the trade-off between the two opposing life history strategies – continued faster growth rates and greater maximum height versus earlier reproduction, curtailed growth, and lower maximum height (Thomas 2011). However, RSOM suffers from the same limitations as maximum height. Thus, both RSOM and maximum height are more usefully seen as outcomes of an RA schedule rather than predictors of it.

While the above mentioned measures of reproductive function may be easier to quantify across large numbers of species, we argue that they cannot substitute for a complete RA schedule. In particular, none of those measures directly captures the instantaneous decision faced by the plant of where to allocate surplus energy, making them difficult to incorporate into process-based models of vegetation dynamics (e.g. Fisher *et al.* 2010; Falster *et al.* 2011; Scheiter, Langan, & Higgins 2013). In contrast, an RA schedule has a direct process-based definition: it specifies the proportion of energy allocated to reproduction as a fraction of the total energy available, at each size or age.

In this paper, our first aim is to review what is known about RA schedules in non-clonal, monoecious, woody plants. Despite several reviews about elements of plant reproduction (Bazzaz *et al.* 2000; Obeso 2002; Moles *et al.* 2004; Weiner *et al.* 2009; Thomas 2011), none have explicitly focussed on RA schedules. This review focuses on perennial species, for recent work has established a framework for investigating RO in annuals, but these methods do not easily extend to perennial plants (Weiner *et al.* 2009). Although studying RA in perennial species is more challenging, these are the species that are the dominant contributors to woody biomass worldwide.

RA schedules show the outcome of a single fundamental trade-off, the allocation of surplus energy to growth versus reproduction, across a plant’s life. As such they summarize some of the most essential elements of a plant’s life history strategy: at what age do plants begin reproducing, what proportion of energy goes to reproduction, and how to plants moderate the proportion of energy they allocate to reproduction as they age. The follow-on information is equally important, for energy not allocated to reproduction is used for growth, increasing the plants height and ability to outcompete its neighbours for light (or nutrients), hence increasing survival. It is, in part, the enormous variation in life history that creates the enormous diversity of plant life forms, and yet we do not know if different RA schedules predict specific life histories or can be predicted by knowing a species’ functional trait values. Here we present a summary of empirical data for the handful of studies quantifying complete RA schedules, as well as some data sets that include only particular features of an RA schedule, such as the shape of the curve. Second, we summarize studies that compared RA or RA schedules across individuals, populations or species growing under different disturbance regimes or with different resource availabilities, and hence give insight on what environmental, life history, or functional traits might alter either instantaneous RA or the RA schedule.

## Methods

### Defining and quantifying reproductive allocation schedules

Figure 4 provides a conceptual outline for the energy budget of a plant, emphasizing that when calculating the amount of energy allocated to growth, it is necessary to distinguish between growth that replaces lost tissues, and growth that increases the size of the plant. Beginning at the box reading “Net production = gross production – respiration”, consider that a plant of a given size and with a given collection of functional traits has a given photosynthetic rate (or gross primary production, GPP) and respiration costs (which consume some of the energy produced, and are subtracted from GPP to yield net primary production, NPP). Some of this plant’s NPP will be used to replace lost or shed tissue, the box labelled “maintenance growth replacing turnover”, with the remainder designated as “surplus energy”. This surplus energy can be allocated to growing to a bigger size or to reproduction, the growth-reproduction trade-off. (Energy can also be allocated to storage or defence, but these are not included in Figure 4.) Note that total growth on the plant in a given year is not one of the boxes, since it represents a combination of energy used to replace lost tissues, i.e. the portion of NPP used to maintain current size, and surplus energy allocated to growing to a bigger size than it was at the start of the survey period.

**Figure 4.**
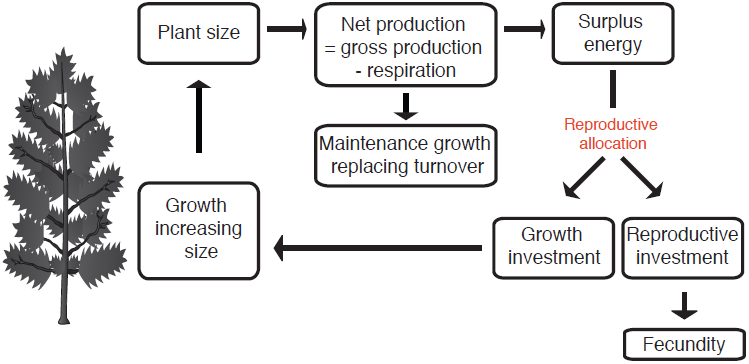
Energy flow within a plant, showing how a given quantity of surplus energy is divided between reproductive investment and growth. Note that total vegetative growth includes maintenance growth, replacing parts lost via tissue turnover, and new growth leading to a net increase in size, termed “growth beyond replacement” in the text.

To properly quantify an RA schedule, one must measure all the energy allocated to growth and reproduction over time. In principle, an RA schedule concerns the instantaneous fraction of surplus energy allocated among growth and reproduction. In practise, the energy budget is typically tabulated on a per year (or per season) basis. The weight of dry biomass is the most commonly used proxy for "energy”, but the kilojoules energy contained in the biomass or the mass of a specific limiting element are valid alternatives. It is important that the same energy units be used for both reproductive and vegetative material.

Reproductive investment should be measured over an entire reproductive cycle and include energy invested both in seed and accessory tissues, the latter termed accessory costs. Accessory costs include the construction of pre-pollination (flower, nectar, and pollen) and post-pollination (packaging, protective and dispersal tissues; aborted ovules) floral parts. Total accessory costs are highly variable and can be as much as 99% or as little as 15% of reproductive energy investment (Table 1).

**Table 1.** Compilation of data from studies measuring reproductive accessory costs. Values give the range of each accessory cost as a percentage, with the mean shown in brackets. Pre-pollination costs are both those required to construct the inflorescence as well as nectar production to entice pollinators, and pollen production. Inflorescence costs include support structures (receptacle, peduncle) and floral parts (sepals, petals, stamens, stigma, ovary, ovules). The post-pollination cost of aborted ovules includes aborted immature seeds at all stages. Packaging, protective and dispersal costs include abiotic dispersal structures, tissue that attracts animal dispersers, and enlarged receptacles. Finally seed cost is the actual cost of the seed, independent of the rest of the fruiting structure.

| authors | species or lifeform | number<br>of<br>species | Pre-pollination costs |  |  | Post-pollination costs |  |  |  |
| --- | --- | --- | --- | --- | --- | --- | --- | --- | --- |
|  |  |  | Inflorescence<br>(%) | nectar<br>production | pollen<br>production<br>(%) | aborted ovules<br>(%) | packaging,<br>protective and<br>dispersal costs<br>(%) | total<br>accessory<br>costs (%) | seed costs<br>(%) |
| Lord & Westoby 2006 | many species and<br>lifeforms | 14 | 0.5-63 (15.7) | not<br>measured | not<br>measured | 0.6-72 (12.9) | 0.7-94 (43.2) | 33.4-96.1 (71.8) | 4-67 (28.2) |
| Henery & Westoby<br>2001 | serotinous Proteaceae | 10 | not measured | --- | --- | included in next<br>category | 90-99 (97.7) | 90-99 (97.7) | 5-55 (2.3) |
| Henery & Westoby<br>2001 | woodland and heathland<br>perennials | 37 | not measured | --- | --- | included in next<br>category | 15-95 (70) | 15-95 (70) | 5-85 (30) |
| Greene & Johnson 1994 | trees | 17 | not measured | --- | --- | data not provided | data not provided | 23-97 (69) | 3-77 (31) |
| Chen *et al.* 2010 | subtropical woody dicots | 62 | not measured | --- | --- | included in next<br>category | 15-98 (47) | 15-98<br>(47) | 2-85 (53) |
| Ashman 1994 | *Sidalcea oregana*,<br>hermaphrodites | 1 | 60 | N/A | 4 | not measured | <1, so ignored | 64 | 36 |

To calculate the investment in growth, one must determine how much bigger the plant is, relative to a year earlier, in units of increased leaf area and increased wood volume. This growth can be tabulated in one of two ways. One can measure both total new vegetative growth as well as leaf, branch and root turnover for one year or develop allometric equations that correlate an easily measured metric, such as diameter at breast height (dbh), with total plant biomass (or another measure of energy investment). A year-on-year increase in dbh then yields the increase in plant size. Surplus energy can also be diverted to storage or the production of defence compounds, two very difficult components to assess. If the stored energy will eventually be used for growth or reproduction, the total lifetime RA will be unaffected by ignoring this energy pool, but the shape of the RA schedule will be skewed.

Unless you are able to follow a single plant through its life, you must find individuals of different sizes, preferably of known age, on which to measure RA. These individuals should be growing under similar environmental conditions and in a similar community of species. One approach to estimating a complete RA schedule for long-lived species, is to pick a known chronosequence, as is available with plantation trees and in locations with a known disturbance (and germination) history (Zammit & Zedler 1993; Cleary, Pendall & Ewers 2008; Genet, Bréda & Dufrêne 2010). Combining RA measurements from plants across a range of sizes yields an RA schedule; a curve showing how an individual’s relative investment in reproduction shifts with plant size or age – i.e. a lifetime’s worth of decisions about energy allocation (Fig. 1). We have focussed on size-related patterns, as size has been shown to have a greater influence on RA than age (Herrera 1991; Pino, Sans & Masalles 2002). In particular, size is the primary factor determining the onset of reproduction in competitive environments (Pino et al. 2002).

### Literature

Here we review what can be learned about RA data from existing studies on 34 populations, representing 32 species. These are the only studies we found in the literature that include data either on how RA changes with size (or age) or that compare RA across populations or closely-related species. We exhaustively searched the literature using both Web of Science and Google Scholar for studies that had measured reproductive investment at multiple ages, across different resource environments or under different disturbance regimes. Some studies used a known chronosequence, some followed the same individuals (or population) across multiple years, and yet others used co-occurring individuals of different sizes to construct a RA schedule. Additional studies report measures of RO, proxies for RA, such as flowering intensity (e.g. Herrera & Jovani 2010) or number of reproductive modules (e.g. Miller *et al.* 2008), but not actual biomass or energy allocation to reproduction. Ideally, RA values were available for individuals at multiple sizes (or ages), such that a RA schedule could be plotted. Knowing RA at reproductive onset and 2-3 later time points is sufficient to predict the shape of the RA schedule, but of course more data points increased the precision with which the RA schedule could be drawn. However, we included studies from which the shape of the RA schedule can be estimated, even if absolute RA values cannot be calculated. The categorization of RA schedule types in Figure 1 is based on a visual assessment, as data are insufficient for a statistical classification. Studies solely reporting plots of reproductive biomass against plant size have not been included as they have been thoroughly reviewed recently (Weiner *et al.* 2009; Thomas 2011) and do not provide any means of determining whether a plant with a large reproductive capacity has a high rate of mass production or large allocation to reproduction. Most of the studies included have not themselves explicitly plotted RA schedules, but instead provide data that can be used to quantify RA schedules (see Appendix 1 for details).

Based on published information, RA was calculated as the proportion of total surplus energy, on a per time basis, allocated to reproduction. One year (or one growing season) is the commonly used time interval. Energy units used are per gram dry mass or kilojoules (determined by burning the samples). Total surplus energy is calculated as the sum of reproductive output (RO), “growth beyond replacement”, as defined in Figure 4, energy stored underground, and energy allocated to defence. RO is the sum total of all types of reproductive investment: flowers, nectar, aborted fruit, mature fruit, and vegetative structures associated only with flowering. Remember, surplus energy does not include growth to replace lost tissue, for tissue replacement costs are subtracted from net primary production to calculate surplus energy (see previous section). It is noted in Table 1 when studies report total new growth, not growth beyond replacement; using total new growth instead of “growth beyond replacement” overestimates surplus energy and underestimates RA. Very few studies consider energy stored underground and energy allocated to defence. When available, these are summed with growth, otherwise this pool is ignored (set to zero). If growth beyond replacement is not directly reported, it is estimated from data on increase in stem diameter and increase in leaf area. RA is then calculated and plotted against plant size (or age) to determine the shape of the RA schedule. Unfortunately, most studies report data for only some reproductive components, usually ignoring shed accessory tissues. The missing reproductive costs are thus not included in our analysis, which will cause RA to be underestimated.

Individual components of an RA schedule are presented in Table 2 and discussed below. They include the shape of the RA schedule, RA at maturation, maximum RA, and size at maturation. For the following papers, the numbers presented in Table 2 were taken directly from the published articles: Pitelka 1977; Pritts & Hancock 1983; Oyama 1990; Alvarez-Buylla & Martinez-Ramos 1992; Comps *et al.* 1994; Ehlers & Olesen 2004; Poorter *et al.* 2005; Read *et al.* 2006, 2008; Miller *et al.* 2008. For the remaining studies, we calculated RA schedules using published data (see Appendix 1 for details).

**Table 2.**
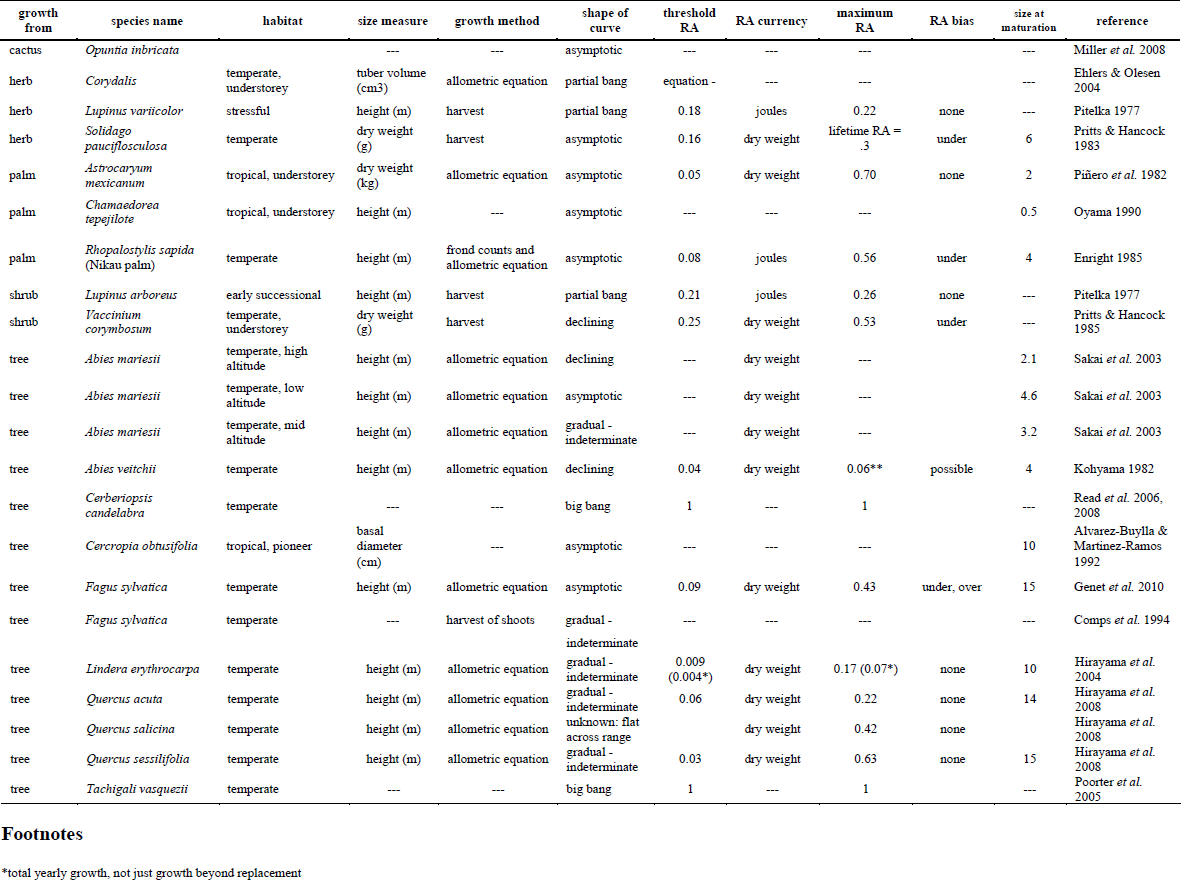
A compilation of available data on reproductive allocation schedules. The shape of the curve is given for all studies, while more precise numbers including RA at the onset of reproduction (threshold RA) and maximum RA are given for the subset of species with available data. The method for determining the plant growth used to calculate RA is given as an allometric equation (derived to correlate a diameter with a specific plant mass) or “harvest” indicating the plants were collected and weighed at the end of the study

## Review of empirical data

### Lifetime reproductive allocation schedule

The species sampled exhibit an enormous variety of reproductive strategies, from truly big bang species (Figure 1, panel b, Table 2) to a great diversity of graded reproduction schedules (Figure 1, panels c-g, Table 2). All but two of the species included exhibit one of the graded RA schedule. Some species, including most perennial herbaceous species studied, ramp up to their maximum RA within a few years of reproductive onset (Pitelka 1977; Ehlers & Olesen 2004) and are classified as “partial bang” (Figure 1, panel b). Other species show a more gradual increase in RA, but still each a definite plateau, the ‘asymptotic’ type in Figure 1, panel d (Piñero, Sarukhan & Alberdi 1982; Oyama 1990; Alvarez-Buylla & Martinez-Ramos 1992; Genet *et al.* 2010). Many of the longest lived species, including both evergreen and deciduous temperate trees, continue to increase RA throughout their lives, never reaching an obvious asymptote (Comps *et al.* 1994; Hirayama, Itoh & Yamakura 2004; Hirayama *et al.* 2008), and are therefore labelled “gradual-indeterminate” (Figure 1, panel e). This collection of RA schedules matches expectations: similar lifetime RO is obtained by a few years of relatively high RA or many years of mostly lower RA. Faster growth allowed a monocarpic species *Tachigali vasquezii* to reach a large size and reproductive maturity more quickly than co-occurring iteroparous species; that is faster growth allowed the onset of reproduction to be advanced (Poorter *et al.* 2005).

In most of the studies considered, the maximum RA achieved is maintained until the end of life, in agreement with evolutionary theory predicting increasing or stable RA until death (Roff 2002; Thomas 2011). However, there are several species, *Vaccinium corymbosum* (Pritts & Hancock 1985), *Abies veitchii* (Kohyama 1982), *Quercus serrata* (Nakashizuka, Takahashi & Kawaguchi 1997) and high elevation populations of *Abies mariesii* (Sakai *et al.* 2003), where RA decreases late in life and thus exhibit a declining RA schedule (Figure 1, panel g, Table 2).

### Reproductive allocation at maturation

Long-lived iteroparous species usually initially have very low RA values, such as 0.05 for *Rhopalostylis sapida* (Nikau Palm) (Enright 1985) and 0.08 for beech (Genet *et al.* 2010) (Table 2). By contrast, shorter lived species can have quite high RA values the year they commence reproduction, such as 0.25 for *Vaccinium corymbosum* (Pritts & Hancock 1985) and 0.18 for *Lupinus variicolor* (Pitelka 1977) (Table 2). Two semelparous perennial species, ones with a big bang schedule where they instantaneously reach RA=1, are included in Table 3. Several hundred additional species are known to have this life history (Young 1984, 2010; Klinkhamer, Kubo & Iwasa 1997; Thomas 2011).

**Table 3.** Studies showing a correlation across populations or closely-related species between RA or threshold size (or age) and a demographic parameter or plant dimensions. The ecological explanation given by the authors is included.

| Study unit | Species | Observed correlation | Ecological explanation | Reference |
| --- | --- | --- | --- | --- |
| Populations | *Attalea speciosa* | Shadier environment -> Larger threshold size | Individuals in lower resource environments must be bigger before they can afford to allocate energy to reproduction. | Barot *et al.* 2005 |
| Populations | *Drosera intermedia* | Higher adult mortality -> Higher RA, in some environments | Individuals with fewer years to reproduce must allocate more energy to reproduction. | De Ridder & Dhondt 1992a; de Ridder & Dhondt 1992b, |
| Species | 4 alpine and subalpine species | Higher elevation (lower resource environment) -> Lower RA | Species in lower resource environments can afford to invest less energy in reproduction. | Hemborg & Karlsson 1998 |
| Species | 3 *Pinguicula* species | Higher adult mortality -> Higher RA | Individuals with fewer years to reproduce must allocate more energy to reproduction. | Karlsson *et al.* 1990; Svensson *et al.* 1993 |
| Half-sib families | *Platago major* | Tall grass environment -> Lower RA | Individuals that were selected to grow taller (in comparison to those in a mown environment), had less energy to invest in reproduction. | Reekie 1998 |
| Populations | *Verbascum thapsus* | Higher mortality -> Smaller threshold size | Individuals in environments that become inhospitable more quickly have fewer years to reproduce and must begin reproducing at smaller sizes. | Reinartz 1984 |
| Populations | *Abies mariesii* | Higher mortality -> Earlier maturation, higher RA | Individuals in environments with greater mortality must begin reproducing earlier and must allocate more energy to reproduction. | Sakai *et al.* 2003 |
| Populations | *Pinus pinaster* | Less favourable environment (PCA of multiple climatic features) -> Higher RA, smaller threshold size (with respect to female function) | Individuals in overall unfavourable environments must begin reproducing earlier and must allocate more energy to reproduction. | Santos-del-Blanco *et al.* 2010, 2012 |
| Populations | *Cynoglossum officinale* | Lower growth rates, higher mortality -> Smaller threshold size | Individuals in overall unfavourable environments must begin reproducing at smaller sizes. | Wesselingh *et al.* 1997 |
| Species | grasses | Poor resource environments --> Lower RA, delayed maturation | Species in lower resource environments must be bigger before they can afford to allocate energy to reproduction and even then allocate less energy to reproduction. | Wilson & Thompson 1989 |

### Maximum reproductive allocation

Maximum RA for semelparous species, such as *Tachigali vasquezii* and *Cerberiopsis candelabra,* is always close to 1 (Poorter *et al.* 2005; Read *et al.* 2006). Iteroparous species usually have a maximum RA between 0.4 and 0.8 (Table 2), although values as low as 0.1 have been recorded in an alpine community (Hemborg & Karlsson 1998). Long-lived iteroparous species are expected to have lower maximum RA than shorter-lived species, as they are diverting more resources to survival, both in the form of more decay and herbivore resistant leaves and stems and other defence measures. These species compensate for a lower RA by having more seasons of reproductive output. However, no clear trend in longevity versus maximum RA is noted among the studies in Table 2, with the highest RA, 0.70, recorded in a temperate palm that lives for more than 250 years.

### Shifts in reproductive allocation with disturbance frequency or resource availability

Comparisons across species or populations that are subject to different environmental conditions have identified certain RA schedule components that recurrently co-vary, suggesting convergent adaptation. In each case, the two populations (or species) grow either in locations that differ in resource availability or in disturbance frequency (effecting mortality), with resultant shifts in RA schedule components. Species or populations with smaller threshold size or earlier maturation, generally have higher RA, supporting traditional life history theory that weedy species have higher fecundity (Stearns 1992; Table 3). Higher mortality is also correlated with this fast-growth strategy, suggesting that the aforementioned traits compensate for having fewer years to reproduce. Lower resource availability is recurrently correlated with lower RA and delayed maturation. Of these studies, only Sakai et al. (2003) have sufficient data to plot complete RA schedules (see Table 3), with the other studies only providing data on portions of the RA schedules such as size at reproductive onset, initial RA or maximum RA.

## Discussion

Using RA schedules to compare reproductive strategies across species (or populations) distinguishes between energy allocated to fundamentally different tissue types and thus links to a key physiological trade-off in an organism’s functioning and life history. Plants that allocate more of their surplus energy to reproduction release more seed but grow less, potentially increasing competition-induced mortality as others that defer reproductive investment progressively overtop the plant. Yet, despite the long-recognised importance of RA (Harper & Ogden 1970), remarkably few RA schedules have been quantified. The limited data available do however suggest that plants display an enormous diversity of RA strategies, ranging from the ‘big bang’ strategy displayed by semelparous species to a variety of graded reproduction strategies, with maximum RA in iteroparous species ranging from 0.21 to 0.71 (Table 2). Moreover, although complete RA schedules are not available, studies that compared RA across populations (or species) with different resource availability or disturbance frequency (Table 3) suggest populations (or species) that are short lived have earlier maturation and rapidly increase RA after maturation. In contrast, lower mortality and later maturation would be associated with a very gradual increase in RA and a slow approach to maximum height (i.e. gradual-indeterminate or asymptotic strategy) This supports demographic data that indicate species growing in higher resource environments exhibit different life history strategies compared to those in low resource environments. Higher resource environments tend to be home to individuals (and populations and species) with low survival, high fecundity, high growth rates, early reproductive maturity, and short life span, versus individuals with the opposite collection of trait values (Bender, Baskin & Baskin 2000; Franco & Silvertown 2004; Forbis & Doak 2004; Garcia, Pico & Ehrlen 2008; Burns *et al.* 2010). As more complete RA schedules become available, it will become possible to refine these patterns and more precisely quantify types of RA schedules (Figure 1) are associated with which life histories. Different functional trait values, including growth rates and energy investment into specific tissues, should also influence RA schedules, but trait-based groupings cannot yet be made, due to limited data availability. In addition, true comparisons of RA schedules can only be made when the same researchers study multiple species with different life history traits, as the many methods used to determine RA likely mean the shapes of the RA curves are more comparable than the values. For now, we can turn to the theoretical literature on optimal energy allocation models, where insights hint at possible factors underpinning interspecific differences.

### Utility of reproductive allocation schedules

Over 40 years ago, Harper and Ogden (1970) recognised the intrinsic value for RA, over RO, in understanding plant function, stating that “Ideally a measure of reproductive effort would involve the determination of starting capital, gross production, and that fraction which is output in the form of propagules.” A focus on the instantaneous allocation of energy by the plant allows RA schedules to be easily incorporated into a variety of process-based plant growth-and ecosystem models (e.g. Fisher *et al.* 2010; Falster *et al.* 2011; Scheiter, Langan, & Higgins 2013). In contrast, knowledge about maximum height, RSOM, or RV curves is substantially harder to integrate, because these trait values provide little information about the partitioning of energy or carbon among different tissue types.

We expect the integration of RA schedules into models would help clarify several empirical patterns. For example, growth rates among larger plants show only weak relationship to leaf and stem traits – this could be because substantial variation in RA among species veils the underlying effects of traits influencing mass production and deployment (Wright *et al.* 2010) If additional empirical data collections of complete RA schedules show that specific functional trait values link with specific RA schedules, these data would allow for improved estimates of seed production after disturbance, a key parameter in management of frequently disturbed vegetation (Noble & Slatyer 1980; Bradstock & Kenny 2003).

Better estimates of instantaneous RA and RA schedules also have the potential to alter estimates of carbon flux and plant growth. For example, in a widely used model of regional carbon uptake and population dynamics, the ecosystem demography model (Moorcroft, Hurtt & Pacala 2001), a fixed fraction (0.3) of surplus carbon is allocated to reproduction. Our results suggest this amount is lower than the maximum achieved by most species, but also that allocation varies substantially through ontogeny.

### What drives variation in reproductive allocation schedules? Insights from optimal energy models

#### Cole’s paradox: A ‘big bang’ strategy is best most of the time

Instantaneous RA implies a clear trade-off between alternative sinks for surplus energy. Starting with this trade-off, optimal energy allocation models identify the RA schedule that maximises seed production across the plant’s lifecycle under a given set of environmental conditions. In the simplest models, surplus energy can only go two places, to “reproductive investment” or “vegetative production beyond replacement” (Figure 4). Optimal energy models that include only this direct linear trade-off find that the complete cessation of growth with reproductive onset, a single reproductive episode and subsequent death (i.e. the big bang strategy from Figure 1, where RA switches instantaneously from 0 to 1) is always optimal, because delayed reproduction and correspondingly greater growth leads to greater final RO (Cole 1954; Kozlowski 1992; Perrin & Sibly 1993; Engen & Saether 1994). Individuals displaying an iteroparous reproductive strategy, with an earlier start to reproduction, an RA less than 1, and multiple reproductive episodes have lower lifetime reproductive output than big bang reproducers. This is because with the iteroparous reproductive strategy, the onset of reproduction leads to decreased growth rates and a smaller adult size, resulting in lower lifetime surplus energy.

In fact, in these models, the reproductive age of big bang reproducers are determined solely by growth rates, how increased size translates to increased reproductive output, and the probability of survival (Kozłowski & Wiegert 1987; Perrin & Sibly 1993); none of these are mechanisms that can be adjusted to create graded RA schedules. This outcome has come to be known as Cole’s Paradox, for the list of perennial semelparous plant species is relatively short, encompassing approximately 100 trees and some palms, yuccas, and giant rosette plants from alpine Africa (see Thomas 2011 for examples), leading researchers on a search for mechanisms favouring a graded reproduction schedule.

#### Non-linear trade-offs or environmental stochasticity are key for promoting graded allocation strategies

One mechanism that generates a diversity of RA schedules is to include non-linear secondary functions in the optimal energy models. The inclusion of a non-linear function may cause the RA schedule to shift away from big-bang to one of the other strategies illustrated in Figure 1.

A large number of optimal energy models investigate how a plethora of factors alter RA (e.g. Myers & Doyle 1983; Sibly, Calow, & Nichols 1985; Reekie & Avila-Sakar 2005; Miller *et al.* 2008). Here we just touch upon some diverse examples that illustrate how non-linear trade-offs lead to graded RA schedules. With a non-linear trade-off, an auxiliary factor causes the cost of increased reproductive investment to increase more (or less) steeply than is predicted from a linear relationship. As a first example, consider a function that describes how efficiently resources allocated to reproduction are converted into seeds. Studying cactus, Miller et al. (2008) showed that floral abortion rates due to insect attack increased linearly with RA. In other words, as RA increases, the cost of creating a seed increases, such that the cacti are selected to have lower RA and earlier reproduction than would be expected from direct costs of reproduction alone. A second example, Iwasa and Cohen’s model (1989) showed that declining photosynthetic rates with size, a trend detected in several empirical studies (Niinemets 2002; Thomas 2010), led to a graded RA schedule. Third, many models, often backed up with data from fish or marine invertebrates, have shown that if mortality decreases with age or size, it benefits an individual to grow for longer and then begin reproducing at a low level – a graded RA schedule (Murphy 1968; Charnov & Schaffer 1973; Reznick & Endler 1982; Kozłowski & Uchmanski 1987; Engen & Saether 1994) (Figure 1, panels c-g). Overall, optimal energy models show that a great diversity of graded RA schedules may be expected, and that as suggested, both fundamental life history traits (mortality, fecundity) and functional trait values (photosynthetic rate, leaf lifespan, growth rates) could affect the shape of the RA schedule. The inclusion of stochastic environmental conditions in optimal energy models also tips the balance in favour of a graded RA schedule (King & Roughgarden 1982).

### Considerations when measuring reproductive allocation schedules

Overall, we advocate for greater measurement of RA schedules. Given RA schedules have been called the measure of greatest interest for life history comparisons (Harper & Ogden 1970; Bazzaz *et al.* 2000), we are surprised by just how little data exist. As described above, we are aware of the variety of challenges that exist to accurately collect this data, including accounting for shed tissue, all reproductive costs, and the yearly increase in size across multiple sizes and/or ages. In addition to these methodological difficulties, we will briefly introduce some other intricacies.

There has been debate as to the appropriate currency for measuring energy allocation. Almost all studies use dry weight or calorie content (joules) as their currency. Ashman (1994), whose study had one of the most complete point measures of RA, showed that carbon content is an inferior predictor of underlying trade-offs compared to nitrogen and phosphorus content, although the general patterns of allocation did not shift with currency. Other studies have found all currencies equally good (Reekie & Bazzaz 1987; Hemborg & Karlsson 1998), supporting the theory that a plant is simultaneously limited by many resources (Chapin *et al.* 1987).

A topic we have not seen discussed in the RA allocation literature is how to account for the transition of sapwood to heartwood. If functionally dead heartwood were considered part of the shed tissue pool, far more of a plant’s annual energy production would be spend “replacing” this lost tissue, decreasing the surplus energy pool and greatly increasing RA for all plants, especially as they approach the end of life. It may even result in more iteroparous species actually approaching RA=1 in old age, as is predicted in many models.

A recent model, however, suggests that reproductive restraint can be beneficial late in life, if it allows an individual to survive for an additional season and have even a few additional offspring (McNamara *et al.* 2009). An alternative hypothesis put forward is that species that can be long-lived may none-the-less benefit from high RA early in life, because the patch environment will be most favourable to the species’ recruitment closer to the time the individual itself germinated (Kohyama 1982; Nakashizuka *et al.* 1997; Ehlers & Olesen 2004). Under this scenario, the species may quickly reach a high RA and later as the patch environment degrades display reproductive restraint if there is a small probability individuals can survive until the patch environment is again ideal for recruitment. This argument most obviously applies to understory species increasingly shaded by a canopy (Pritts & Hancock 1985; Ehlers & Olesen 2004), but was also proposed by Kohyama (1982) to explain decreasing RA with stand age in a canopy tree. Alternatively these patterns may result from incomplete measurements, such as underestimating tissue turnover rates (Figure 4). At this point, there is just too little data to draw many general conclusions, or assess whether methods of data collection are influencing our results.

### Future directions

Two of the most important applications of RA schedules are to provide the necessary data to parameterise and validate vegetation models and to test predictions about changes needed to generate a variety of RA schedules. To make better comparisons and determine more generalities, data for RA schedules must be collected across many species using identical methods. Life history and functional traits must be measured for each species in order to determine how variation in these traits correlates with consistent differences in RA schedules. For instance, such data will allow inter-specific correlations to test whether certain RA schedule parameters are correlated and how the patterns match expectations of life history theory. For decades, theoreticians have been using RA schedules as a fundamental evolvable trait (Iwasa & Cohen 1989). It’s time we empiricists provided some data.

## Acknowledgments

Funding from the Australian Research Council to D Falster and M Westoby supported this work. We thank J Camac, J Johansson and F Thomas for comments on earlier versions of this paper, and M. Henery for access to data used to produce Fig 3.

## 9. Appendix 1

For the studies where RA was not directly reported, we calculated the values as follows:

**Enright 1985.** The manuscript provides data on net annual production across all tree sizes, dividing a plant into leaves, stem, underground tissues, and reproductive tissues. Total leaf number is constant among reproductively mature plants, and therefore new leaf production is simply replacing shed tissues. RA is therefore calculated as yearly reproductive production divided by the sum of net annual production of stems, underground tissues, and reproduction. True RA values will be somewhat higher than those reported, especially for the oldest plants, as root turnover rates are not known.

**Genet 2010.** The manuscript provides data on net annual increase in stem biomass, annual non-structural carbohydrate production, and seed production. Reproductive biomass includes seed and cupule mass, but not other accessory costs. RA is calculated as seed production divided by the sum of increase in stem biomass, annual non-structural carbohydrate production, and seed production. Data were collected in a significant mast year for *Fagus sylvatica*, such that RA averaged over several years would likely be lower. For *Fagus*, the accessory costs ignored should be small compared with seed and cupule mass, but their inclusion would increase RA.

**Hirayama 2004.** The manuscript provides data on increase in “woody organ” biomass, total leaf biomass, and reproductive costs. Reproductive costs include accessory costs, including the number of flowers, aborted fruit, and mature fruit. The allometric equation they used to determine increase in “woody organ” biomass and leaf production are based on averages for an entire community and are not specific to the species they studied. They do not have data on plant age or yearly increase in leaf biomass, such that it is difficult to estimate proportion of total leaf biomass that replaces shed tissue and proportion that represents an increase in leaf biomass. As a rough approximate, we used average annual increase in dbh across all tree sizes (as there was no clear trend with tree size) to estimate difference in age between their 5 trees. We then divided the difference in leaf weight between trees by the difference in age to determine the annual increase in leaf mass. This rough calculation indicated that the majority of leaf biomass replaces the previous year’s shed leaves, while a small fraction represents an increase in leaf biomass. RA is then calculated as reproductive production divided by a sum of increased wood biomass, increased leaf biomass, and reproductive production. This yields RA values up to 3 times greater than presented in their manuscript, although the pattern, a continued increase in RA with height is unchanged.

**Hirayama 2008.** The manuscript provides data on increase in woody material, increase in leaf biomass, and reproductive investment. Reproductive investment includes mature fruit (including dispersal and protective material) as well as aborted flowers and fruits. RA is calculated as reproductive investment divided by the sum of increase in woody material, increase in leaf biomass, and reproductive investment. These species show a distinct pattern of biennial masting, so we averaged RA during a high and low mast year to determine the trees’ actual allocation patterns. Note that the manuscript uses total leaf biomass, not incremental increase, in their calculation of RA, such that their estimates of RA are much lower than the ones we have used. The manuscript does not provide a threshold size or age for these species, only indicating they used a range of dbh values above 20 cm. *Quercus salicina* has a constant RA across this range, leading us to believe the smallest trees included are larger than trees at the “threshold size”. We therefore do not include a threshold size, age, or RA for this species.

**Kohyama 1982.** The data are from a figure in the manuscript. The manuscript does not indicate how “net reproductive effort” is calculated so we are uncertain whether it accounts for shed tissues. The reproductive investment component includes accessory tissues associated with the cone and seed formation. Masting occurs every 4 years and numbers presented in the paper are averaged across years to determine the long-term RA.

**Pinero 1982.** The manuscript includes data on net annual production of roots, trunk, live leaves, dead leaves (interpreted to mean shed leaves), seeds, and reproductive accessory tissues. RA is calculated as the sum of reproductive production divided by the sum of net annual production of roots, trunk, live leaves and reproductive materials.

**Pritts 1985.** The manuscript includes data on annual biomass production of vegetative and reproductive structures for a range of plant ages (their Figure 3). Vegetative structures are divided into stems, roots, and leaves. Total new vegetative production is the sum of new stem production, new root production, and the increase in leaf biomass over the previous year. We use the increase in leaf production as *Vaccinium corymbosum* is a winter deciduous species and the other portion of leaf production offsets previously shed tissue. RA is calculated as reproductive production divided by the sum of reproductive and new vegetative production.

**Sakai 2003.** The manuscript provides equations for increment increase of wood (R^2^H increment) and annual reproductive biomass for three populations of *Abies mariesii*. Neither annual leaf and root production, nor turnover of these structures, is known. RA could therefore not be calculated, although the possible shape of the RA curve could be. Using the equations provided and estimating a wood density of 0.5, annual wood production and reproductive production are determined. RA is then calculated as reproductive production divided by the sum of wood production and reproduction production. If a hypothetical increase in leaf area is included, shifting from double wood production in young plants to a small fraction of wood production in mature plants, the shapes of the curves of the middle and low population plants are unchanged, while the initial RA at the high site is much reduced and the RA across plant size is fairly unchanged.

